# Draft genome sequence of a predatory bacterium from northern peatland soil

**DOI:** 10.1101/2025.10.08.681201

**Authors:** Tatiana Demina, Riina Ihonen, Minna K. Männistö, Jenni Hultman

## Abstract

Predatory bacteria are abundant in soil, but their diversity and functions remain not fully understood, especially in subarctic regions. Here, we report strain 1-FT3.2, a predatory bacterium obtained from peatland soil in Northern Finland (Pallas, 68 °N). The bacterium was cultivated on *Mucilaginibacter cryoferens* FT3.2 as prey. Although a pure culture of strain 1-FT3.2 was not obtained, its draft genome was assembled from sequencing reads derived from the co-culture with its prey. The draft genome of 1-FT3.2 is 7.2 Mb in length and 81% complete. Genome analyses suggested that 1-FT3.2 belongs to the family *Polyangiaceae* (phylum *Myxococcota*), which comprises predatory bacteria. The genome annotation revealed (near-)complete metabolic modules of central carbon metabolism and aerobic respiration. Two proviral regions were predicted in the draft genome, both putatively representing tailed phages of the class *Caudoviricetes*. Several CRISPR-Cas system proteins were also identified. The draft genome sequence could be used in future comparative studies assessing the diversity of predatory bacteria in northern soils or other environments.

## Introduction

Predatory bacteria are important players in microbial food webs (Hungate et al. 2021). Myxobacteria are a group of bacteria associated with the phylum *Myxococcota*, characterised by group predatory behaviour and a complex lifestyle, where rod-shaped vegetative cells can aggregate into multicellular fruiting bodies and produce spores (Saggu et al. 2023). Myxobacteria are globally distributed and especially abundant in soil (Zhou et al. 2014; Wang et al. 2021). Together with other micropredators, myxobacteria play leading roles in carbon sequestration and mineralization in soil (Lueders et al. 2006). Moreover, myxobacteria may dominate among other potential bacterivores and have been suggested to represent one of the keystone taxa in soil microbial food webs (Petters et al. 2021). Still, more data are needed to resolve their taxonomic diversity as well as metabolic and lifestyle capacities across environments, including relatively underexplored subarctic regions.

Since soil microbial communities are highly diverse, obtaining complete genomes through metagenomics may be a challenging task (Anthony et al. 2024). Cultivating soil microbes makes it possible to reconstruct their genome sequences reliably and link genetic information to the observed phenotype. In this study, we obtained strain 1-FT3.2, a predatory bacterium from northern peatland soil in the Pallas region, Finland, using *Mucilaginibacter cryoferens* FT3.2 (Kumar et al. 2025) as prey. *M. cryoferens*, recently described as a new species, was isolated from Arctic tundra soils in the Kilpisjärvi region, Finland, where it may play important roles in litter decomposition and carbon recycling together with other *Mucilaginibacter* species (Männistö et al. 2009; Kumar et al. 2025). Strain 1-FT3.2 remained in a mixed culture with its prey, but the analyses of its draft genome sequence obtained from the co-culture suggest that it belongs to the *Polyangiaceae* family.

## Methods

### Soil sampling, isolation and cultivation conditions

A soil sample was collected from peatland in the Pallas area, Northern Finland, in September 2022 (N67°59’ E24°13’, Figure 1A). The vegetation was mainly sedges (Figure 1B). The sample was collected from a depth of 5 cm with sterile instruments and stored at 4°C. The pure culture of *Mucilaginibacter cryoferens* FT3.2 (Kumar et al. 2025), was used as the prey for isolating predatory bacteria from the soil sample. Bacteria were cultivated using R2A medium (Neogen), which contained 0.5 g L^-1^ yeast extract, 0.5 g L^-1^ meat peptone, 0.5 g L^-1^ casamino acid, 0.5 g L^-1^ glucose, and 0.5 g L^-1^ starch and was adjusted to pH 6. For solid and top agar, 15 and 4 g L^-1^ of agar (Sigma-Aldrich) were added, respectively. The cultures were grown aerobically at room temperature (RT).

**Figure 1.**
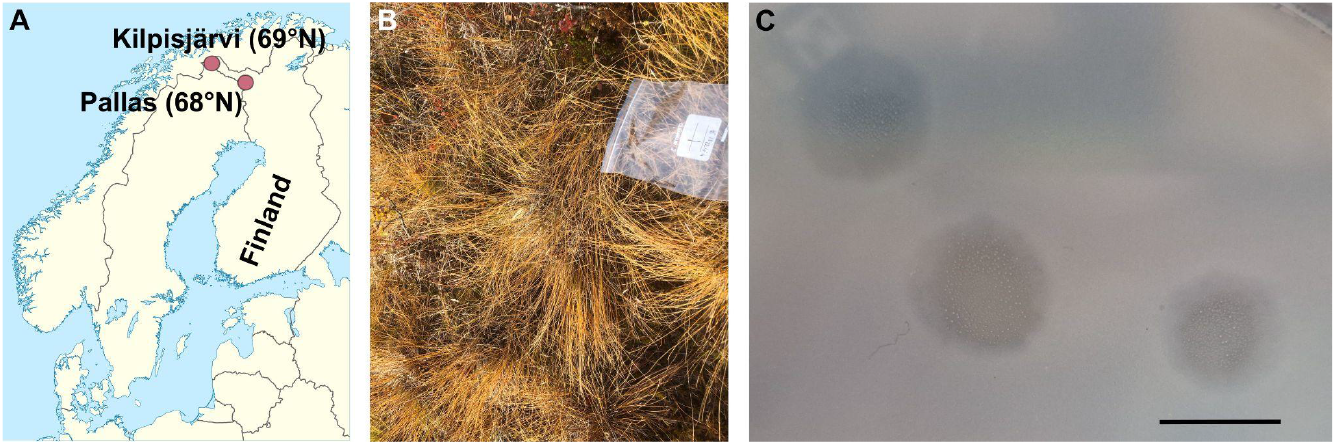
(A, B) Sampling location, Pallas. In (A), additionally, Kilpisjärvi, the original isolation location for the prey, *Mucilaginibacter cryoferens* FT3.2, is shown. Map modified from Wikimedia Commons (NordNordWest). (C) A representative plate with lysis zones on the *M. cryoferens* FT3.2 lawn after 14 days of incubation, scale bar, 1 cm.

For the isolation, 5 g of the soil sample (wet weight) was resuspended in 50 ml of R2A broth and incubated on a shaker (200 rpm) at RT for two weeks for the sample enrichment. The enriched sample was centrifuged (ThermoScientific F15-6×100y, 30 min, 2,500 g, 20°C) and 100 μl of non-diluted supernatant plated with 300 μl of the *M. cryoferens* FT3.2 liquid culture and 3 ml of R2A soft agar (46°C) as a top layer on R2A solid agar plates. The plates were incubated aerobically at RT. The observed growth inhibition/lysis zone was picked up with a sterile pipette tip, resuspended in R2A broth, and plated in a top agar layer as before, which was repeated three consecutive times.

### DNA extraction and sequencing

The top agar layers of the semi-confluent plates were collected and resuspended in R2A broth (3 ml per plate), incubated with shaking (∼200 rpm) at RT for one hour and centrifuged (ThermoScientific F15-6×100y, 30 min, 10,000 g, 4°C). The supernatant was collected and stored at 4°C. The stock titers were determined by plating serial dilutions in a top agar layer as described above. DNA was extracted with the GeneJET Genomic DNA Purification Kit (Thermo Scientific) using the manufacturer’s protocol for Gram-negative bacteria and 20 ml of the agar stock as input. Note that the agar stocks contained cells from both *M. cryoferens* FT3.2 and the new strain.

For sequencing, 100 ng of genomic DNA was converted to a sequencing library using the Illumina DNA prep. Samples were dual indexed using the sequencing core unit’s own Nextera primers. Seven cycles were used in the PCR step and DNA was pooled and purified using Illumina’s SPB bead purification. The Library pool was sequenced at 12 pM on the AVITI sequencer (Element Biosciences) using the AVITI 2×150 Sequencing kit Cloudbreak FreeStyle High Output. Sequencing was performed at the DNA Sequencing and Genomics Laboratory (supported by HiLIFE and Biocenter Finland funding), Institute of Biotechnology, University of Helsinki.

### Genome annotation

FastQC v. 0.11.9 (https://www.bioinformatics.babraham.ac.uk/projects/fastqc/) was used to assess the quality of reads. Raw reads were trimmed and adaptors removed with Cutadapt v. 2.7 (-m 50 --nextseq-trim 20) (Martin 2011). Read-based taxonomic profiling was performed using PhyloFlash v. 3.4.2 and SILVA138.1.eukmod database (Gruber-Vodicka et al. 2020). Since the sample contained a mixed culture of the prey strain *M. cryoferens* FT3.2 and a new potentially predatory strain, SPAdes v. 3.15.5 in the --meta mode was used for genome assembly (Bankevich et al. 2012). BBTools Stats was used for assessing the assembly statistics, Reformat for sorting scaffolds by their GC content, and Dedupe for dereplicating scaffolds (minidentity=95 absorbrc=t absorbmatch=t sort=length) (sourceforge.net/projects/bbmap/). The full-length SSU rRNA gene sequences obtained from the PhyloFlash run and the assembled scaffolds of ≥10 kbp in length were searched with BLASTN (Altschul et al. 1990) against the NCBI nt database using an E-value cutoff of 0.001.The quality of the draft genome of a new strain was assessed with CheckM2 v. 1.0.1 (Chklovski et al. 2023), and GTDB-Tk v. 2.3.2 with GTDB release 226 database (Chaumeil et al. 2022) was used for assigning a taxonomic classification. For the genome annotation, DRAM v. 0.1.2 (Shaffer et al. 2020) was used at KBase (Arkin et al. 2018). Putative (pro)viral sequences were predicted by geNomad v. 1.7 (Camargo et al. 2023) and their quality and completeness assessed with CheckV v. 0.8.1 (Nayfach et al. 2021). Bowtie2 v. 2.5.3 was used for the additional mapping of reads to putative viral sequences (Langmead and Salzberg 2012).

## Results

### Isolation

After about two weeks of incubating the plates, growth inhibition/lysis areas of 4-5 mm were observed. In subsequent platings, the size of lytic zones reached up to about 1 cm (Figure 1C). The central parts of these zones were clear, while edges were hazier. Agar stocks produced lysis zones on the *M. cryoferens* FT3.2 lawn when diluted up to 10000-fold, but no lysis zones could be observed when titrating filtered stocks (0.22 and 0.45 μm PES LLG-Syringe filters Spheros), suggesting that the origin of the observed lytic zones was not viral. Very small, almost transparent or whitish colonies growing over the lysis zones were observed (Figure 1C), but no aggregated structures like fruiting bodies were seen. Despite our attempts, these tiny colonies could not be transferred to a fresh plate for independent growth. An alternative cultivation approach using the myxobacterium-suited CY-C10 medium ((Karwowski et al. 1996) modified by omitting antibiotics) and higher incubation temperature (28°C) for stock titration did not improve colony growth visibility. We named the strain causing lytic zones on *M. cryoferens* FT3.2 as 1-FT3.2.

### Genome sequencing and assembly

Sequencing genomic DNA of a mixed culture resulted in 245,936,278 raw read pairs (150 bp + 150 bp), of which 245,436,350 pairs were retained after read trimming. With the read-based profiling by PhyloFlash, 225,532 reads (0.092% of all reads) could be mapped to SSU rRNA sequences in SILVA database. Of the mapped reads, 212,996 (94%) were assigned to the order *Sphingobacteriales* (*Bacteroidota*), where the genus *Mucilaginibacter* belongs to, and 9,050 (4%) were assigned to the order *Polyangiales* (*Myxococcota*). The rest of the hits constituted less than 0.01% of mapped reads each. Thus, read-based profiling suggested two strains present in the sample, comprising about 98% of reads together. Furthermore, full-length SSU rRNA gene sequences assembled by SPAdes, matched to SILVA database, were only two OTUs with the closest-matching references of *Mucilaginibacter* sp. M20-56 (*Sphingobacteriales*; GenBank acc. no.: KP899210.1, 99% id., 100% cov.) and *Phaselicystis* metagenome (*Polyangiales*; GenBank acc. no.: FPLS01001412.1, 95% id., 99% cov.). Additional BLASTN searches of the two detected OTUs against the NCBI nt database resulted in hits to 16S rRNA gene sequences of *Mucilaginibacter* sp. strain FT3.2 (100% id., 100% cov., 0 E-value) and the members of the order *Polyangiales* (the genera *Minicystis, Sorangium, Chondromyces, Labilithrix, Polyangium*, and uncultured bacterium, 91-92% id., 100% cov., 0 E-value), respectively.

The assembly of the mixed culture consisted of 6,318 scaffolds, of which 140 scaffolds were longer than 10 kbp and represented 95% of the total length of all scaffolds (Table 1). Most scaffolds longer than 10 kb were characterised by a GC content of either 41-43% (71 scaffolds) or 64-66% (56 scaffolds) (Figure 2). The *Mucilaginibacter cryoferens* FT3.2 genome GC content is known to be 42.1 % (Genbank acc. no. CP183228.1). Therefore, 56 scaffolds with a GC content of 64-66% were separated from the rest of the assembly, representing strain 1-FT3.2. Dedupe run confirmed the non-redundancy of the assembled draft genome. The total length of the 1-FT3.2 draft genome was 7,202,438 bp with the scaffolds ranging from 13,622 to 664,534 bp (Table 2). Based on the CheckM2 assessment, the genome is 81.3% complete and 0.5% contaminated.

**Table 1.** Statistics for the mixed culture assembly, listed as of different minimal scaffold length thresholds.

| Minimum scaffold length, bp | Number of scaffolds | Number of contigs | Total scaffold length, bp | Total contig length, bp | Scaffold contig coverage, % |
| --- | --- | --- | --- | --- | --- |
| 50 | 6,318 | 6,369 | 15,040,305 | 15,035,385 | 99.97 |
| 100 | 1,456 | 1,507 | 14,717,700 | 14,712,780 | 99.97 |
| 250 | 299 | 350 | 14,552,681 | 14,547,761 | 99.97 |
| 500 | 222 | 271 | 14,529,291 | 14,524,391 | 99.97 |
| 1,000 | 201 | 250 | 14,513,761 | 14,508,861 | 99.97 |
| 2,500 | 168 | 217 | 14,459,058 | 14,454,158 | 99.97 |
| 5,000 | 155 | 203 | 14,409,144 | 14,404,344 | 99.97 |
| 10,000 | 140 | 188 | 14,302,883 | 14,298,083 | 99.97 |
| 25,000 | 116 | 161 | 13,895,326 | 13,890,826 | 99.97 |
| 50,000 | 93 | 136 | 13,006,064 | 13,001,764 | 99.97 |
| 100,000 | 56 | 89 | 10,405,941 | 10,402,641 | 99.97 |
| 250,000 | 6 | 12 | 2,322,273 | 2,321,673 | 99.97 |
| 500,000 | 1 | 3 | 664,534 | 664,334 | 99.97 |

**Table 2.**
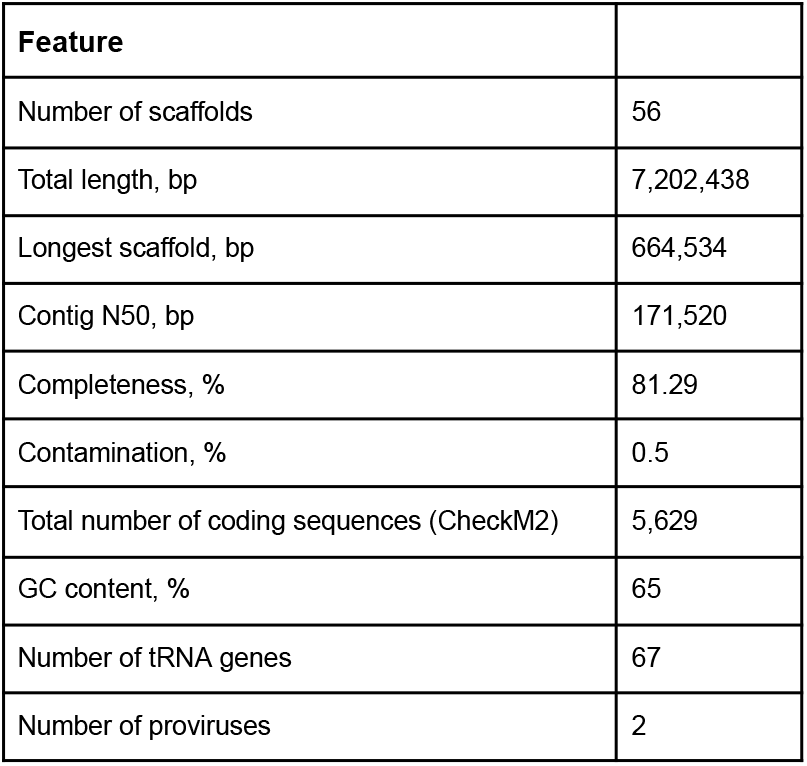
1-FT3.2 draft genome features.

**Figure 2.**
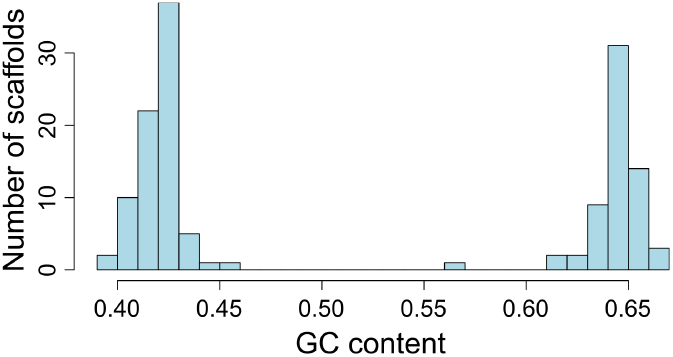
The distribution of GC content across assembled scaffolds longer than 10 kbp.

### Genome classification and annotation

In the BLASTN search, the 1-FT3.2 draft genome scaffolds recruited numerous hits to sequences representing the phylum *Myxococcota*. GTDB-Tk run on the draft genome suggested classifying 1-FT3.2 within the family *Polyangiaceae*, order *Polyangiales*, class *Polyangia*, phylum *Myxococcota*. With DRAM, no rRNA encoding genes were identified in the draft genome scaffolds. DRAM-based annotations (Figure 3) revealed a few complete metabolic modules: pentose phosphate cycle, citrate cycle (TCA cycle), glyoxylate cycle, cytochrome c oxidase, and F-type ATPase, as well as a near-complete (8/9) glycolysis module, suggesting robust central carbon metabolism and aerobic respiration. Also, arsenate reductase (glutaredoxin), acetyl-CoA synthetase, acetate kinase, and alcohol dehydrogenase were predicted, but no CAZy enzymes. The incomplete nature of the draft genome sequence precludes full understanding of metabolic capacities or the lack of those in 1-FT3.2. Among other DRAM predictions, several different CRISPR-Cas system proteins were identified (Cas1, Cas2, Cas3, CasA, CasB, CasC, CasD, CasE, Cmr1, Cmr2, Cmr3, Cmr4, Cmr5, and Cmr6). About 36% of all predicted proteins had no significant hits to any DRAM database.

**Figure 3.**
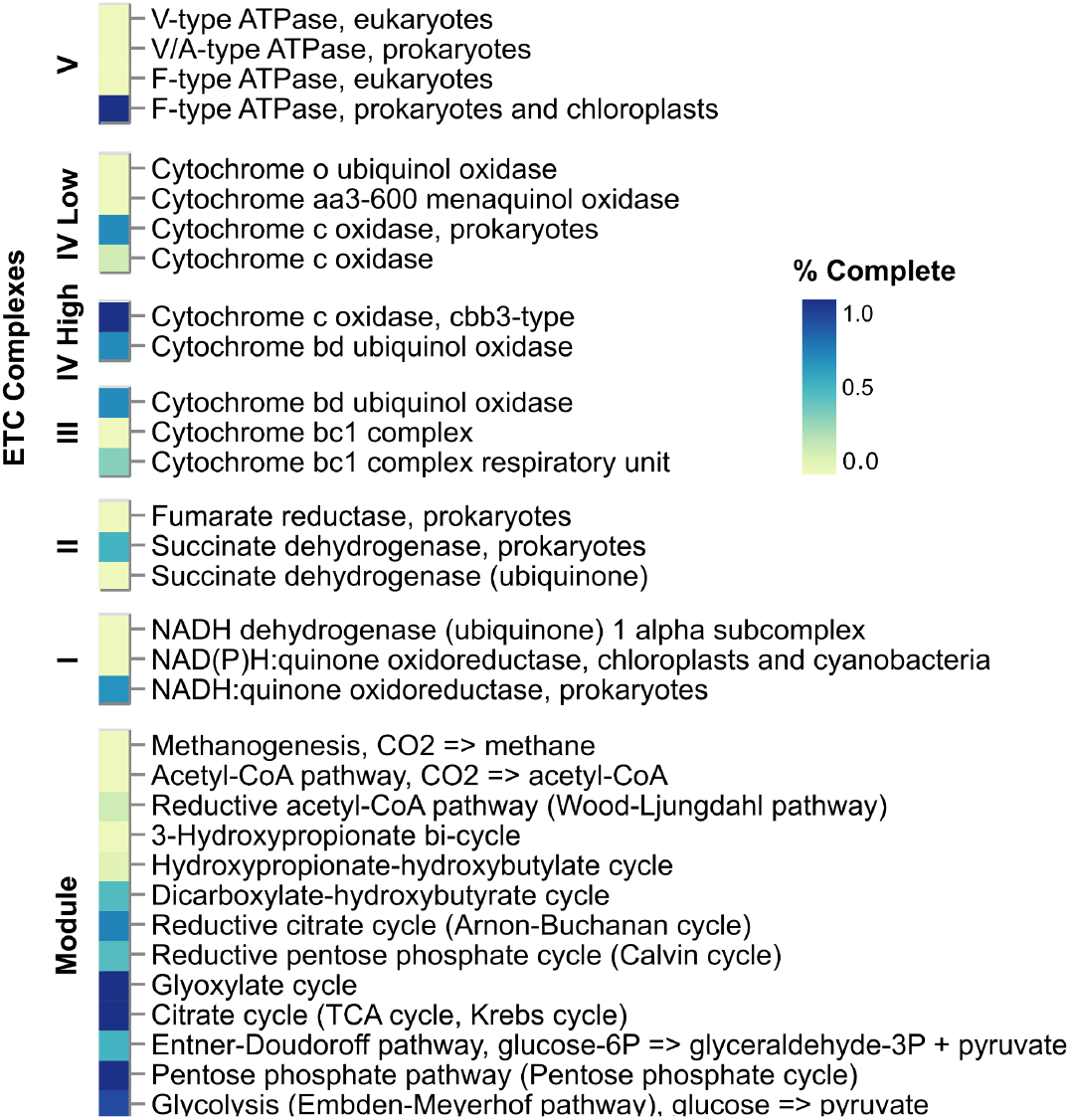
Metabolic functions of 1-FT3.2 strain predicted with DRAM.

Using geNomad with all scaffolds from the mixed-culture assembly resulted in the prediction of two proviral sequences on scaffolds that belonged to the 1-FT3.2 draft genome: at coordinates 58-30,107 nt in NODE_10_length_239568_cov_144.087670 and 23,906-79,669 nt in NODE_67_length_79671_cov_165.578627. These proviral elements were medium-quality (80 and 53 % complete, respectively) and both assigned as tailed phages within the class *Caudoviricetes*. In addition, three other short scaffolds (0.2, 5.4, and 6.9 kbp), were identified as viral by geNomad, although the presence of viral genes could be confirmed by CheckV only for one of them. Mapping reads to these three short scaffolds resulted in an overall alignment rate of only 0.00002%, confirming that the nature of the observed lysis zones is unlikely to be viral.

## Conclusions

The genome analysis of 1-FT3.2, the new predatory bacterium strain reported here, placed it within the family *Polyangiaceae* (*Myxococcota*). Members of this family are terrestrial isolates mainly from soil and plant decay material, characterised by large genomes and high GC content, with some strains being able to degrade cellulose and produce various secondary metabolites (Garcia and Müller 2014). *Polyangiaceae* representatives are rarely isolated from subarctic soils (Dawid 2000). The draft genome sequence and genome of 1-FT3.2 could be used in future comparative studies aiming to resolve the diversity of the family *Polyangiaceae* and/or more broadly, predatory bacteria residing in subarctic soils. Although the reported genome is incomplete, it still contributes to increasing the sequenced space of the soil microbiome. Having the strain available for future laboratory studies makes it possible to explore its lifestyle and metabolic capacities in more detail.

## Data availability

Raw reads from the mixed culture are available from NCBI’s Short Read Archive (SRA): PRJNA1337162. The new strain 1-FT3.2 draft genome is available from Figshare: https://doi.org/10.6084/m9.figshare.30277690.v1.

## Acknowledgements

We thank Erin Way and Essi Suomilammi for technical assistance. We acknowledge DNA Sequencing and Genomics Laboratory (supported by HiLIFE and Biocenter Finland funding), Institute of Biotechnology, University of Helsinki for sequencing and CSC – IT Center for Science, Finland, for computational resources as well as for technical support. This work is supported as part of the Genomic Sciences Program DOE Systems Biology Knowledgebase (KBase) funded by the U.S. Department of Energy, Office of Science, Office of Biological and Environmental Research under Award Numbers DE-AC02-05CH11231, DE-AC02-06CH11357, DE-AC05-00OR22725, and DE-AC02-98CH10886.

